# Sepsis Reconsidered: Identifying novel metrics for behavioral landscape characterization with a high-performance computing implementation of an agent-based model

**DOI:** 10.1101/141804

**Authors:** Chase Cockrell, Gary An

## Abstract

Objectives: Sepsis affects nearly 1 million people in the United States per year, has a mortality rate of 28–50m% and requires more than $20 billion a year in hospital costs. Over a quarter century of research has not yielded a single reliable diagnostic test or a directed therapeutic agent for sepsis. Central to this insufficiency is the fact that sepsis remains a clinical/physiological diagnosis representing a multitude of molecularly heterogeneous pathological trajectories. Advances in computational capabilities offered by High Performance Computing (HPC) platforms call for an evolution in the investigation of sepsis to attempt to define the boundaries of traditional research (bench, clinical and computational) through the use of computational proxy models. We present a novel investigatory and analytical approach, derived from how HPC resources and simulation are used in the physical sciences, to identify the epistemic boundary conditions of the study of clinical sepsis via the use of a proxy agent-based model of systemic inflammation. Design: Current predictive models for sepsis use correlative methods are limited by patient heterogeneity and data sparseness. We address this issue by using an HPC version of a system-level validated agent-based model of sepsis, the Innate Immune Response ABM (IIRBM), as a proxy system in order to identify boundary conditions for the possible behavioral space for sepsis. We then apply advanced analysis derived from the study of Random Dynamical Systems (RDS) to identify novel means for characterizing system behavior and providing insight into the tractability of traditional investigatory methods. Results: The behavior space of the IIRABM was examined by simulating over 70 million sepsis patients for up to 90 days for the following parameters: cardio-respiratory-metabolic resilience; microbial invasiveness; microbial toxigenesis; and degree of nosocomial exposure. In addition to using established methods for describing parameter space, we developed two novel methods for characterizing the behavior of a RDS: Probabilistic Basins of Attraction (PBoA) and Stochastic Trajectory Analysis (STA). Computationally generated behavioral landscapes demonstrated attractor structures around stochastic regions of behavior that could be described in a complementary fashion through use of PBoA and STA. The stochasticity of the boundaries of the attractors highlights the challenge for correlative attempts to characterize and classify clinical sepsis. Conclusions: HPC simulations of models like the IIRABM can be used to generate approximations of the behavior space of sepsis to both establish “boundaries of futility” with respect to existing investigatory approaches and apply system engineering principles to investigate the general dynamic properties of sepsis to provide a pathway for developing control strategies. The issues that bedevil the study and treatment of sepsis, namely clinical data sparseness and inadequate experimental sampling of system behavior space, are fundamental to nearly all biomedical research, manifesting in the “Crisis of Reproducibility” at all levels. HPC-augmented simulation-based research offers an investigatory strategy more consistent with that seen in the physical sciences (which combine experiment, theory and simulation), and an opportunity to utilize the leading advances in HPC, namely deep machine learning and evolutionary computing, to form the basis of an iterative scientific process to meet the full promise of Precision Medicine (right drug, right patient, right time).

## Introduction

There are nearly 1 million cases of sepsis in the United States each year, with a mortality rate between 28–50% [1] While operational care process improvements in the last 20 years that have led to reduction in mortality [2] therapeutic options for sepsis remain variations of anti-microbial and physiological support dating back nearly a quarter century, and, crucially, there remains no FDA approved biologically targeted therapeutic for the treatment of sepsis. In an era where the overall goal of medical care is to provide personalized/precision medicine, which should mean “right drug for the right patient at the right time,” achieving this goal in sepsis is significantly hampered by the lack of success in translating basic science knowledge into robust and effective therapeutics - a problem pervasive across the biomedical spectrum [3]. Over a quarter century of failed attempts to modulate sepsis raises a key question: how tractable are current investigatory strategies, namely increasing granular clinical data collection and analysis coupled with pre-clinical experimental investigations, aimed at achieving this goal? Is there a fundamental limitation of the capabilities of the current approaches to attempting to characterize and control sepsis? This question of the epistemic limits of traditional research methodology is not limited to sepsis, but is arising across the biomedical research spectrum. For example, a recent paper examines such methodological limits by applying current analytical approaches used in neuroscience to a known, simpler proxy information processing system, a microprocessor, and finds those approaches insufficient for meaningful characterization of the behavior of the microprocessor [4]. We propose that a similar approach can be used to evaluate the tractability of increasing reductionist attempts to define and characterize clinical sepsis. We use a known computational model, a previously developed agent-based model of sepsis recognized to be vastly simpler than the real world system, and apply to it the types of data extraction and system state characterization possible with respect to clinical biomarker analysis. This use of computational/simulation models as proxy systems has a long history in systems engineering, and has been described specifically in the use of biomedical agent-based models (ABMs) [5]. The rationale for using a simulation proxy model to establish analytical boundary conditions is based on the fact that there is complete knowledge of the computational model, i.e. there are no “hidden variables,” and therefore it has an internally consistent ground truth. As a computer program, every component and interaction of the model is known, including probabilities associated with stochastic events inputted into the model’s code. Given that the ABM is vastly simpler than the real world system, we assert that if complete knowledge of the behavioral output of the simpler proxy model is insufficient to meaningfully predict its behavior, attempts to do so in the real world will be similarly futile. We propose that an iterative process utilizing large-scale simulation of sufficiently complex proxy models and advanced analysis of simulation data can establish “boundaries of futility” with respect to what can or cannot be known about clinical sepsis utilizing data-centric methods. This approach is only made possible by the exponential growth of computing power seen in leadership-class high performance computing (HPC) that now makes tractable the ability to near comprehensively characterize the behavioral landscape of an ABM. We utilize an HPC implementation of a previously developed system/population-level validated ABM of the innate immune response [6] (the Innate Immune Response ABM or IIRABM) to generate a first approximation behavioral landscape of sepsis to guide the development of suitable metrics with which to characterize that behavioral landscape. Though developed over 15 years ago, the IIRABM remains valid insomuch it reproduces the overall dynamics of the acute inflammatory response, producing recognizable system level outcomes (healing, pro-inflammatory death, hypo-immune death and overwhelming initial infection) using a knowledge-based rule system that, while admittedly incomplete, has not been demonstrated to be incorrect in its represented behaviors by accumulated discoveries since its development. In fact, some of the behaviors generated by the IIRABM presaged their general recognition within the sepsis community, specifically the temporal concurrence of pro- and anti-inflammatory cytokine responses (as opposed to sequential pro- and compensatory responses) [7,8] and the importance of the immuno-incompetent/attenuated recovery phase of sepsis, particularly with respect to its prolonged duration [9–11]. For the current investigations, we acknowledge the incompleteness of the IIRABM, considering it an abstract, substantially less complex representation of the human innate immune response. As such, we further pose that the multi-dimensional behavioral landscape of the IIRABM is less complex than can be expected of the real world system, and that characterization of the behavioral landscape of the IIRABM represents a lower bound, first approximation of the ability to characterize the real world system. The investigations presented here are divided into two distinct but related tasks: 1) identification of the region of parameter space of the IIRABM that corresponds to biologically/clinically plausible behavior, and 2) the development of novel metrics and dynamical systems analysis to characterize the behavior of the IIRABM and provide insight into the tractability and viability of current pathways toward precision medicine for sepsis.

## Materials and Methods

For our investigations we use a previously validated agent-based model (ABM) of sepsis, the Innate Immune Response ABM (IIRABM) [6]. The IIRABM was ported from NetLogo [12] to C++ and implemented using MPI (Message Passing Interface) 2.0 for the parallelization and distribution of the model. The IIRABM utilizes cells as its agent level. Individuals are represented as an interaction surface between endothelial cells and circulating inflammatory cells and are subjected to variable perturbations representing either infectious insult or tissue trauma. Multiple cell types are represented, including endothelial cells, macrophages, neutrophils, TH0, TH1, and TH2 cells, as well as precursor cells for these cell lines. Details for the IIRABM can be seen in Ref [6]. System damage is represented by an aggregate measure of individual endothelial cell damage; the death threshold is set at 80%, representing the ability of ICU care to render short-term lung, kidney, and cardiac failure to be survivable [6]. The increasing death threshold is important for two reasons: it simulates ICU care in that modern medical techniques allow a human to survive an amount of damage that would not have been survivable before modern medicine; and an increasing death threshold allows the model to evolve to states which would have been unreachable if execution were halted earlier [6,13]. Beyond the components and interactions of the inflammatory response, the IIRABM utilizes 4 “external variables” that describe clinical features that affect, but are not systemically integral to, the clinically-relevant systemic inflammation: environmental toxicity (recurrent microbial exposure seen in a clinical environment), host resilience (an aggregate variable representing the cardiorespiratory reserve of the patient and manifest as the ability of damaged tissue to recover its oxygen level), and two measures of microbial virulence, invasiveness (ability to spread in host tissue) and toxigenesis (ability to kill host tissue) (See Supplemental Table 1). Note that these variables represent factors that clearly influence an individual’s response during sepsis, but that there are currently no clinically accessible, or even identifiable metrics that would allow categorization of a particular patient. It is because of this combination of known influence and inability to categorize that the entire parameter space needs to be characterized. In total, 8800 environmental parameter sets were represented. A set of infectious injuries, the consequences of which ranged from trivial to severe, was applied to each of these parameter sets. This injury was represented by a circular region with a radius that varied from 1 to 40 grid-spaces in diameter. The inherent stochasticity built into the IIRABM represents both the real/biological and epistemic variability seen in the immune response to injury and infection; this allows the model to capture the variability seen in a clinical population. Thus, identical initial injuries can lead to diverging patient trajectories depending on how the immune system of an in *silico* patient responds to the simulated pathogen and injury distribution. The primary basis for the behavioral validity of the IIRABM is its ability to reproduce the general dynamics of a system’s inflammatory response to infectious insult resulting in four clinically relevant trajectories: healing, hyper-inflammatory system death, immuno-paralyzed late death, and overwhelming infection [6]. These validity criteria are conserved from the initial description of the IIRABM, and are based on the common sense observation that a biologically plausible system should not be always killed by the smallest insult possible, nor should it always completely recover from the largest insult possible. Therefore, a biologically plausible parameter set for the model would have a lower threshold of system injury from which it would always recover (i.e. “healed”) as well as an upper threshold of system injury from which it would always die (i.e. “overwhelming infection). The additional dynamic classes of behavior become evident within this zone of plausible behavior, and arise due to the fundamental purposes of the inflammatory response, namely the eradication of infection by invasive microbes (enhanced by forward feedback processes to increase its efficacy) and facilitation of the recovery from system damage (which mandates negative feedback control to attenuate the initial response). Both of these dynamics result in the effective eradication of invading microbes, but result in subsequent system death due to the disordered internal processes. While each of these dynamic classes represents a different type of control failure (pro-inflammatory death = distorted forward feedback and immune-paralysis = distorted negative feedback) for purposes of this current initial behavioral analysis we classify them together as representing internal system failure, clinically analogous to multi-organ failure. Future analysis will separate these two dynamics, but within this current paper we define three possible system level outcomes: 1) healing, 2) system failure and 3) overwhelming infection. We use these three criteria as the basis of population distribution characterization for the simulated individual cohorts. The work presented in this paper describes the outcomes of an initial parameter sweep consisting of 70.4 million simulations (8800 parameter sets × 40 injury sizes × 100 stochastic replicates × 2 treatments). Subsequent simulation experiments were performed to refine the development of system-describing metrics and behavioral properties of the IIRABM as described below. Simulations were run on Edison Cray XC30 Supercomputer at the National Energy Research Scientific Computing Center and on Beagle Cray XE6 Supercomputer at the University of Chicago. The vast data output of these simulations required the development of novel means of characterizing and visualizing data at multiple-scales of analysis; it is not technically feasible to represent this information in a single form. Therefore, we have developed as set of nested metrics to emphasize specific perspectives aimed at establishing a baseline behavioral landscape of clinical sepsis. We utilize five primary metrics with which to characterize the behavior of the system: 1) population-level outcome distributions for each parameter set 2) multi-dimensional parameter space characterization across the entire range of parameter sets, 3) severity of outcome distribution heatmaps to reify across biologically plausible parameter sets 4) probabilistic basins of attraction (PBoA) and 5) stochastic trajectory analysis. The first 3 metrics are used to identify the region of parameter space of the IIRABM that corresponds to biologically/clinically plausible behavior and confirm the baseline validity of the IIRABM.

## Results

We evaluated the behavioral space of the IIRABM with respect to 10 values of toxigenesis, 4 values of invasiveness, 20 values of host resilience, and 11 values of environmental toxicity (See Supplemental Table 1). Each combination of these parameters was evaluated with N = 100 stochastic replicates for each initial injury represented as a circular injury with a radius ranging from 1 to 40 cell widths.

### 0.1 Metric #1. Population-Level Outcome Distributions

The results of the simulations (N = 100) for each parameter set shown as a histogram with injury size as the x-axis, and the count of a specific outcome for the y-axis and is used to examine outcome-severity progression for an individual parameter set as a function of the magnitude of the injury simulated. The three possible outcomes of the IIRABM — healing, system failure (sepsis), or overwhelming infection, are represented using three possible bars for each injury number. This approach was initially described in rudimentary form in [6] and forms the basis by which the biologically/clinically plausible overall parameter space characterization is performed. For the metrics listed below, Metric #2 can be considered to utilize the collapse of Metric #1 into a 0-dimensional point data representation, whereas Metric #3 can be considered to utilize the collapse of Metric #1 into a 1-dimensional line representation. Figure 1 demonstrates 3 representative population outcome distributions as a function of injury size. Where Figure 1A depicts a non-plausible condition where the system always dies with infection and Figure 1B depicts a non-plausible condition where the system always recovers from infection, Figure 1C depicts a plausible, clinically relevant condition where at some level the system heals, at some level the system dies from infection, and in between there is the generation of behavior where the system clears the infection yet dies for system failure (i.e. most of clinical sepsis). This parameter set had moderate invasiveness (= 2) and toxigenesis (= 4) values, low environmental toxicity (=1), and average host resilience (= 0.9). With an injury radius of 22 cells, approximately 50% of the *in silico* patients lived and 50% died due to system failure. We consider the population outcome distribution seen in 1C as an example of a clinically relevant parameter set, since all 3 classes of outcomes (healing, system failure and overwhelming infection) are present.

**Figure 1.**
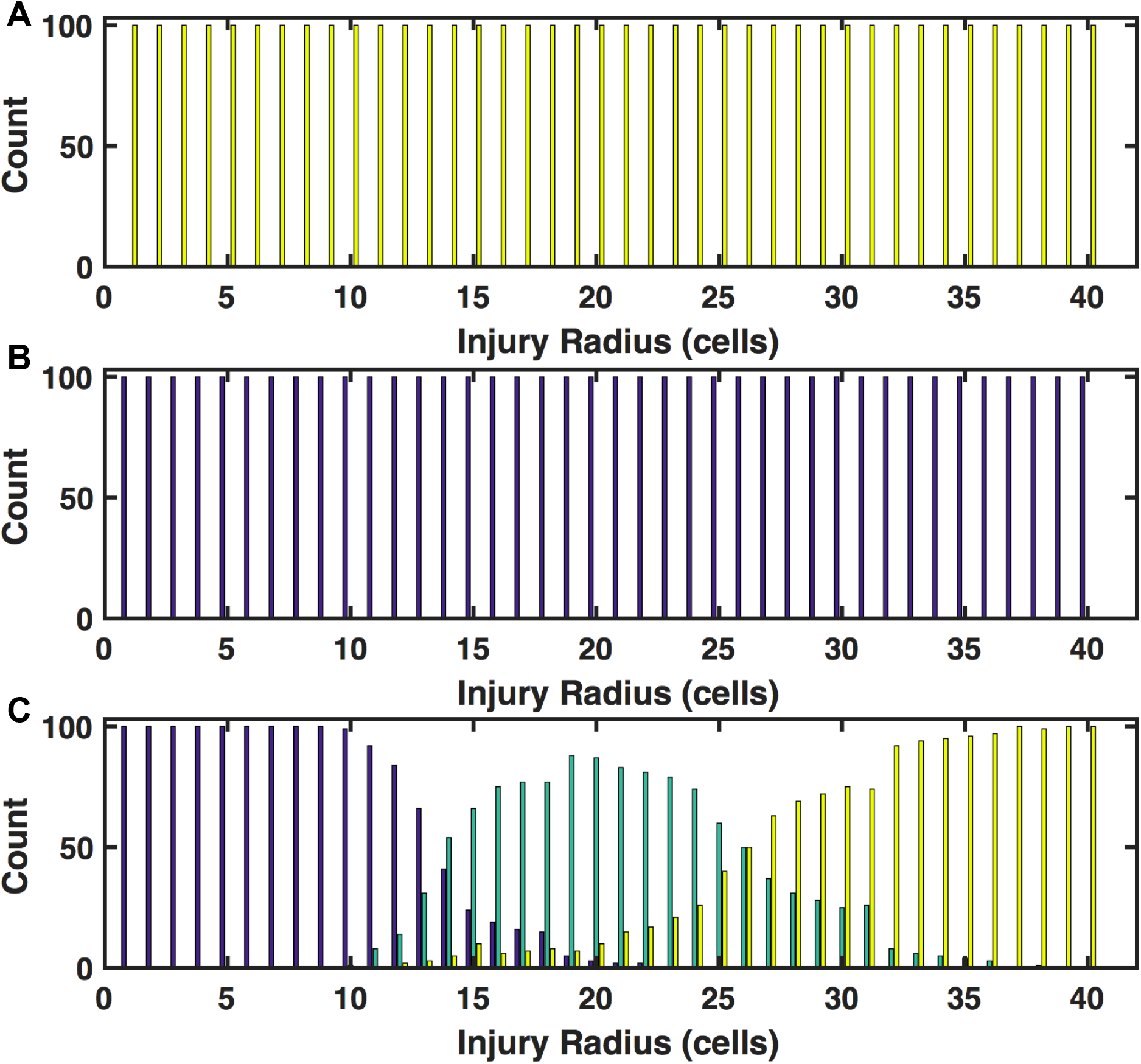
Population Outcome Distributions across a range of initial infection numbers. Outcome counts for a specific injury size are shown on the y-axis. Blue shading indicates complete healing, green shading indicates death by sepsis, and yellow shading indicates death by infection. Panel A displays a histogram of outcomes as a function of injury size for simulations run with high bacterial virulence (invasiveness=4, toxigenesis=10), low host resilience (resilience=0.05), and high environmental toxicity (environmental toxicity=10); this parameter set is implausible because the system always dies with even the minimal injury. Panel B displays a histogram of outcomes as a function of injury size for simulations run with low bacterial virulence (invasiveness=1, toxigenesis=1), high host resilience (resilience=0.9), and low environmental toxicity (toxicity=1); this parameter set is implausible because the system always heals despite the maximal injury. In Panel C, the full range of outcomes as a function of initial injury size is displayed for an infection with moderate toxigenesis (toxigenesis=4) and invasiveness (invasiveness=2), low environmental toxicity (environmental toxicity=1), and average host resilience (resilience=0.1). Smaller initial injuries (shown on the left of the figure) more often result in complete healing of the simulated host (as expected) while the largest initial injury values, which also deliver the highest infectious load, result in death from overwhelming infection. The central region of this panel shows a region in which the host is capable of fighting off the applied infection, but is unable to recover from the inflammatory state that was required to fight the infection.

### 0.2 Metric #2. Multi-dimensional Parameter Space Characterization

A multi-dimensional parameter sweep across a range of initial perturbation (initial infection) identifies the upper and lower bounds with respect to clinically plausible behavior, as reflected in Metric #1 above. The lower bound is that defined by parameter combinations where the system cannot be killed by the minimal, least virulent microbes (i.e. always heals); the upper bound is defined by parameter combinations where the system cannot recover from the maximal, most virulent microbes (always overwhelmed by infection). Therefore, biologically plausible/clinically relevant parameter space resides between these two boundaries. The representation of this parameter space is a multi-dimensional grid, where each point in the grid represents a classification of the outcomes with respect to Metric #1; points in that space that reflect biologically plausible behavior is then termed Clinically Relevant space. The distribution of population outcome distributions across the entire parameter space investigated can be seen in Figure 2. Each 3-axis graph represents the configuration of parameter space with respect to toxigenesis, invasiveness and host resilience; because of the inability to represent 4 axes, different levels of environmental toxicity are represented by different panels. All parameter space representations demonstrated a qualitatively consistent structure with plausible boundaries reflecting the range of possible human behavior in response to microbial sepsis. Panels A and B display outcome spaces for untreated patients with low environmental toxicity (toxicity=1) and high environmental toxicity (toxicity=10) respectively. Panels C and D display outcome spaces for patients treated with antibiotics with low environmental toxicity (toxicity=1) and high environmental toxicity (toxicity=10) respectively. Each point represents 4000 in *silico* patients (40 injury sizes, each with 100 stochastic replicates). Points are color-coded based on the outcomes generated. Blue points represent simulations that healed under all circumstances. These points are distributed in regions of space where host resilience is high and the bacterial virulence is low (lower invasiveness and lower toxigenesis). Red points represent simulations that always died from overwhelming infection; these points are distributed in regions of high bacterial virulence (higher values for invasiveness and toxigenesis). Black points represent simulations that either died from overwhelming infection or healed completely and mark the boundary between simulations that always heal and simulations always die from infection. Pink points represent simulations which either died from overwhelming infection or hyperinflammatory system failure; these points are found primarily in the simulations that were treated with antibiotics and had low values for environmental toxicity and host resilience. Globally, the IIRABM behaves as expected; more virulent bacterial infections combined with less resilient hosts more often lead to a negative outcome while a combination of less virulent bacteria and more resilient hosts is more likely to lead to complete healing. Furthermore, the application of antibiotics shifted the outcome space such that the overall probability of death by sepsis or infection was globally reduced (Pink Points in Figure 2). The central region of highest outcome uncertainty, in which the simulation can heal completely, die from overwhelming infection, or die from system failure (SIRS/MOF), corresponds to the critically ill population (Green Points). This region of parameter space defines the Clinically Relevant space, producing the most diverse model dynamics and as such can provide the greatest insight into the fundamental structure of the model. As previously noted, there is currently no way of knowing where individual real world patients would reside within the Clinically Relevant space: while all 4 of the external variables represent “real” factors determining the severity of sepsis, there is no current way of even qualitatively correlating those rankings with any clinically accessible metric. This means that there is no way to know what the overall distribution is of all sepsis patients across the Clinically Relevant space, and that any attempt to “match” a mortality rate from a particular parameter set to that seen in clinical sepsis (i.e. the 20–50% mortality generally reported [1,2]) would be a false and misleading attempt at demonstrating “validation.”

**Figure 2.**
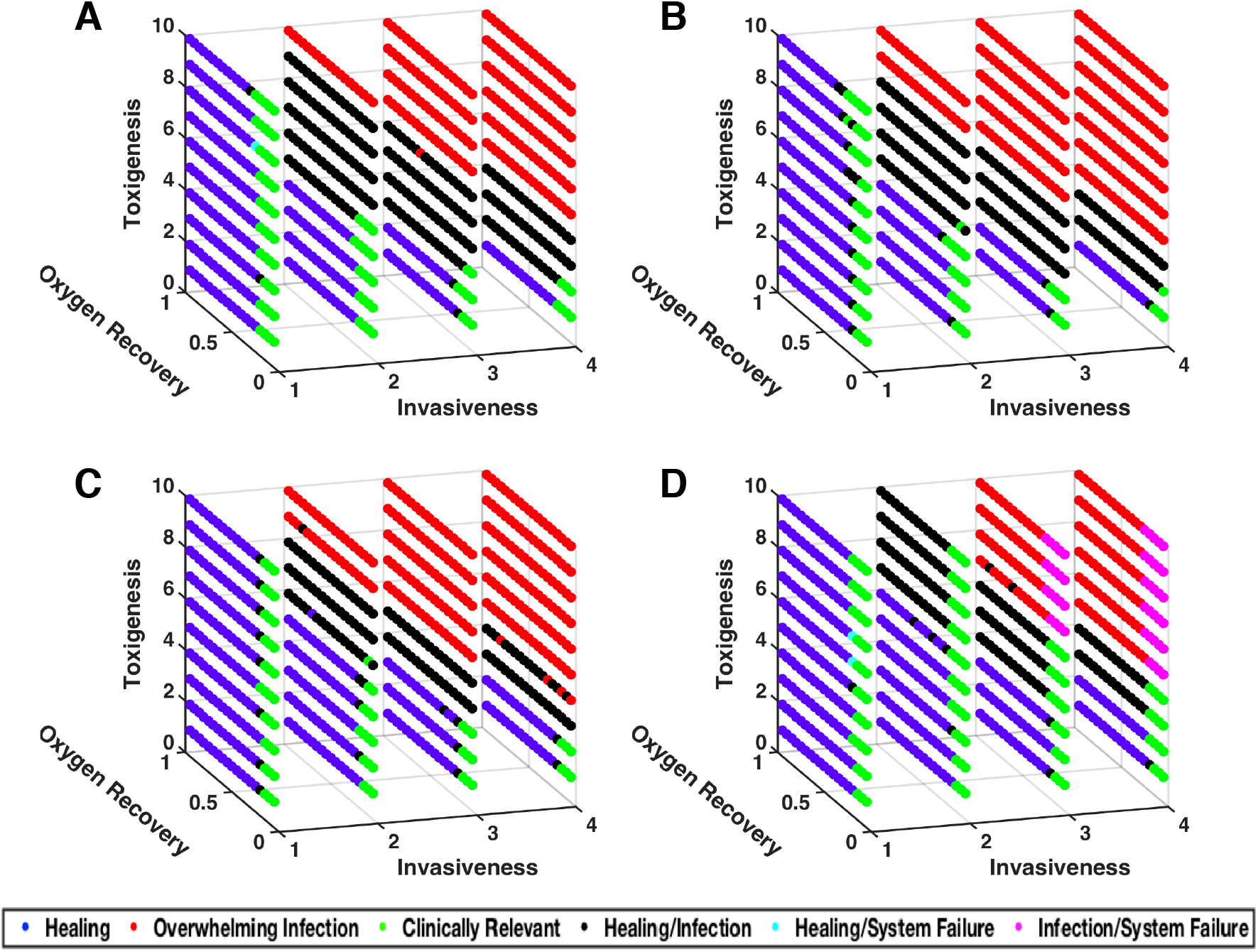
Visualization of outcome space for sepsis model. Panels A and B display outcome spaces for untreated patients with low environmental toxicity (toxicity=1) and high environmental toxicity (toxicity=10) respectively. Panels C and D display outcome spaces for patients treated with antibiotics with low environmental toxicity (toxicity=1) and high environmental toxicity (toxicity=10) respectively. Each point represents 4000 *in silico* patients (40 injury sizes, each with 100 stochastic replicates). Points are color-coded based on the outcomes generated. Blue points represent simulations that healed under all circumstances. These points are distributed in regions of space where host resilience is high and the bacterial virulence is low (lower invasiveness and lower toxigenesis). Red points represent simulations that always died from overwhelming infection; these points are distributed in regions of high bacterial virulence (higher values for invasiveness and toxigenesis). Black points represent simulations that either died from overwhelming infection or healed completely and mark the boundary between simulations that always heal and simulations always die from infection. Pink points represent simulations which either died from overwhelming infection or hyperinflammatory system failure; these points are found primarily in the simulations that were treated with antibiotics and had low values for environmental toxicity and host resilience. Green points represent the Clinically Relevant simulations as these parameter sets lead to all possible outcomes; these points are distributed in regions of low to middle values of the host resilience parameter and moderately virulent infections. For all classes of simulation, the final outcomes are primarily dependent on the host resilience and microbial virulence parameters and have a secondary dependence on environmental toxicity.

### 0.3 Metric #3. Severity of Outcome Distribution Heatmaps

Though all parameter sets in the CR space can generate all of the three possible model outcomes, there is still significant variation in their behavioral distributions. It is therefore desirable to be able to reify the parameter sets by their outcome distributions in order to visualize the configuration of severity across the entire relevant portion of parameter space. Severity of outcome distribution heatmaps present a simplified view of outcome space for a chosen group of parameter sets, i.e. a condensed form of representing the population distributions seen in Metric #1. Rather than view each element in a parameter set as an individual axis, we assign a ranking to each parameter set (based on the binary outcome of mortality). The set of these rankings then makes up the Y-axis. We use the X-axis to display the range of injury sizes to which an individual parameter set is exposed ala Metric #1. The outcome distribution (percentage of *in silico* patients that ultimately died for a given parameter set and injury size) is then represented by the colors of the heatmap. The representation of a 1-dimensional heatmap of the population outcome distribution allows the reification of initial parameter sets based on the robustness/fragility of the particular parameter set across biologically/clinically plausible parameter space, and the response of the overall system to a putative therapy (i.e. antibiotic therapy). These visualizations help confirm the plausible structure of the population distributions of the simulated experiments. In Figure 3, outcomes are ordered from “worst” on the top of the figure to “best” on the bottom of the figure. In this case, worst is defined as the parameter set for which death instances within the population begin at the smallest injury size. Each horizontal line in both panels represents 4000 simulations of a single parameter set. The shading indicates the percentage of deaths; blue represents no deaths at and yellow represents 100% death. Figure 3A and Figure 3B show identical parameter sets but different ordering, where patients represented in the Figure 3B received antibiotics twice a day for 10 days beginning 6 hours after their injury. As expected, the application of antibiotics greatly increases the survivability of the simulated injuries – the yellow region is moved to the right, indicating improved survival to higher initial injury numbers when antibiotics are available. The antibiotic parameter set was reordered (again from worst to best) as the application of antibiotics changed the system dynamics. The “stepping” feature that can be seen in Figure 3B is due to an emergent clustering of parameter sets by the host-resilience parameter. The most resilient hosts are on the top and the least resilient are on the bottom of each step group. This makes sense because the application of antibiotics significantly decreases the bacterial infection’s influence on the system; the other external parameters, invasiveness, toxigenesis, and environmental toxicity, are all directly related to the simulated microbial infection. When antibiotics are applied, the host-side parameters take precedence for outcome determination, a plausible behavior further substantiating the validity of the IIRABM. Following the use of these first 3 metrics to identify the biologically plausible Clinically Relevant regions of parameter space for the model, we then turn to the development of 2 novel data analysis methods in order to evaluate basic properties of the system and how they might impact the investigation of real-world, clinical sepsis.

**Figure 3.**
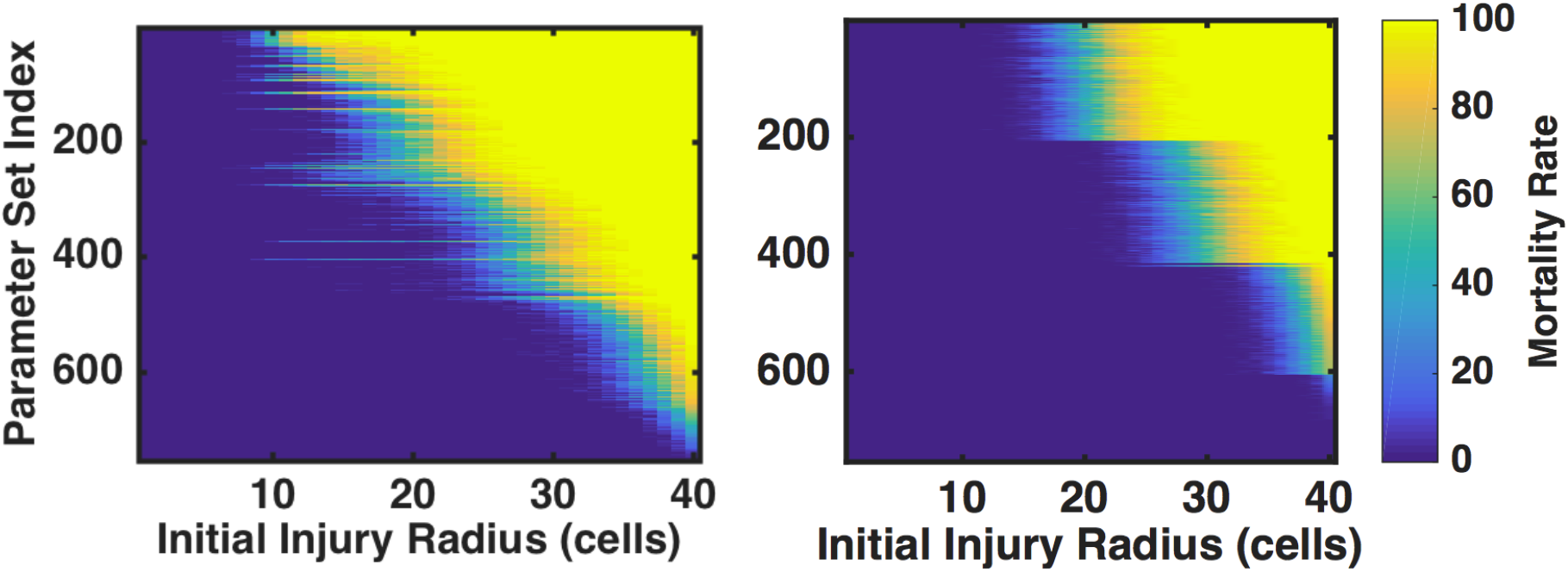
Heatmaps indicating percentage of death as a function of initial injury size for clinically interesting parameter sets. Patients either went untreated (left) or received antibiotics twice/day for 10 days beginning 6 hours after their injury. Shading indicates the percentage of simulations that lived or died; areas that are dark blue represent complete healing 100% of the time, and areas shaded yellow represent death (either from infection or sepsis) 100% of the time. Note that each row of this figures is essentially a compressed version of Figure 1. Parameter sets are ordered from worst (meaning highest chance of dying at the lowest injury number) on the top to best on the bottom. There is no clear structural relationship between parameter sets for the untreated case (left), though the simulations treated with antibiotics self-stratify based on the host-resilience parameter (with lower host resilience parameters found towards the top of the figure). This stratification is due to the semi-neutralization of microbial virulence through the application of antibiotics.

### 0.4 Metric #4. Probabilistic Basins of Attraction (PBoA)

The current approach to disease classification/prognosis/diagnosis is to use some set of measurements (be they biomarkers or physiological signals) taken from the patient at a particular point in time, with the goal that such information, or time series sequences of such information, will be able to distinguish between groups with some threshold of acceptable success. The more general description of this concept is characterizing the relationship between system state and system trajectory, the basis of the study of Dynamical Systems [14] – that is, a system that evolves in time according to some rules in which future states evolve from the current state. In dynamical systems theory, an attractor is a state or set of states to which certain states tend to evolve [15]. A basin of attraction is the region of system state space under the influence of a given attractor as it affects given trajectories passing through these points (think analogously of gravitational wells in astronomy). The stochasticity inherent to the IIRABM, and the real world system as well, makes it a Random Dynamical System (RDS) [16] as random events influence the system’s evolution. Because this system is not deterministic, a true basin of attraction cannot be mathematically defined. However the concept is still conceptually valid with the addition of a “stochastic zone” where the direction of a particular trajectory is uncertain. When this model is viewed as an RDS, it becomes clear that there are two attractors, one leading to an oxygen deficit of 0 or complete healing (referred to as the Life Attractor), and one leading to an oxygen deficit of 8160 or death (referred to as the Death Attractor), with a stochastic zone of uncertain trajectory direction in between. To characterize the multi-dimensional attractor space for the IIRABM as a RDS, we have developed a metric termed the Probabilistic Basin of Attraction (PBoA). The PBoA represents a map of possible states within parameter space available to the model. At each point, the probability that the system evolves to a specific attractor (i.e. system-level outcome) is calculated from the results of a large number of simulations. The PBoA map generated can be used to qualify the interplay between outcomes and measured variables. The full PBoA is a high-dimensional object with one axis for each measured variable; for instance, this model generates a 20-dimensional PBoA. As it is difficult to display such a high-dimensional object, we show 2-dimensional projections of the PBoA along axes of interest. The configuration of the PBoA for a particular set of system measurements will give insight into the utility of those measurements as predictors of system outcome. The PBoA also addresses the currently stated limits to predictive modeling by providing near comprehensive state space coverage (thereby addressing the data paucity limitation) for a system where all components are known (thereby addressing the “hidden” variable limitation). We focus on the Clinically Relevant portion of parameter space to identify the attractor space across the multiple variables present in the IIRABM, with the goal being to determine the feasibility of using individual or sets of potential biomarkers as predictors of system outcome. We present 2-dimensional “slices” of IIRABM PBoAs in Fig. 4, with the output of 3,016,000 simulations represented (those within the Clinically Relevant zone). The blue dots represent circumstances where all simulations recovered, the crimson dots represent conditions where all simulations died, with the intervening heatmap depicting the progressive probability of system death (= the stochastic zone). Figure 4, Panels A and B depict the predictive capability of values for IL-10 and TNF respectively across a range of values of system damage. The large and irregularly shaped stochastic zone without a clear structure across a range of system damage suggests that a particular level of either of these cytokines alone poorly correlates with system outcome and would be an inadequate biomarker for outcome (this is consistent with the current understanding of the utility of IL-10 or TNF as a marker of disease severity). Figure 4C shows the probability of death as a joint function of TNF and IL10 levels. Interestingly, the “slope” of the region of greatest mortality is not a constant, but levels off as the concentration of IL10 increases. This suggests that an intervention augmenting systemic IL10 production would be insufficient to significantly alter a patient’s outcome or disease trajectory. Figure 4D, demonstrating the PBoA for population levels of TH1 cells versus system damage, shows a qualitatively similar structure as to Figure 4A and B: there appears to be virtually no independent predictive capability for the number of TH1 cells present. Figure 4E displays the PBoA for population levels of TH2 cells against system damage, where the “slope” of the region of highest mortality has a slight upward slope, suggesting that higher levels of TH2 cells can be associated with negative outcomes in sepsis. However, the width of the vertical distribution of outcomes is large enough to wash out the predictive capability of this trend. Alternatively, plotting the population of TH1 cells versus the population of TH2 cells (Figure 4F) generates a more interesting PBoA that demonstrates the benefits of being able to shift between multi-dimensional data projections. Here we see a very poorly bounded stochastic zone for lower levels of both T-cell subtypes, likely representing the earlier phases of the disease course as the system evolves, that transitions into a different region of state space with a more defined structure. In this region the predictive capability of the TH1/TH2 ratio becomes more powerful, with an increase in the proportion of TH2 cells associated with adverse outcomes, a finding that is consistent with both existing reports [17] and the negative consequences of later immunoparalysis [9–11]. However, in general, the extent/breadth/width of stochastic zones within the PBoAs point to the insufficiency of snap shots of system state as a predictor of system outcome. We confirm this finding by performing principle component analysis (PCA) on the Clinically Relevant region of the IIRABM using total system state capture at intervals of 6 hours for the first 24 hours post infection, and then every 24 hours for up to 5 days; this sequence was chosen to encompass the scope of current investigations into prognostic biomarkers for sepsis [18,19]. These PCA plots (Supplemental Material 1) show no discrimination at any of the time points, consistent with the findings of the PBoAs. Given the predictive insufficiency of system state snapshots, we now turn to more directly evaluating the trajectory space of the IIRABM.

**Figure 4.**
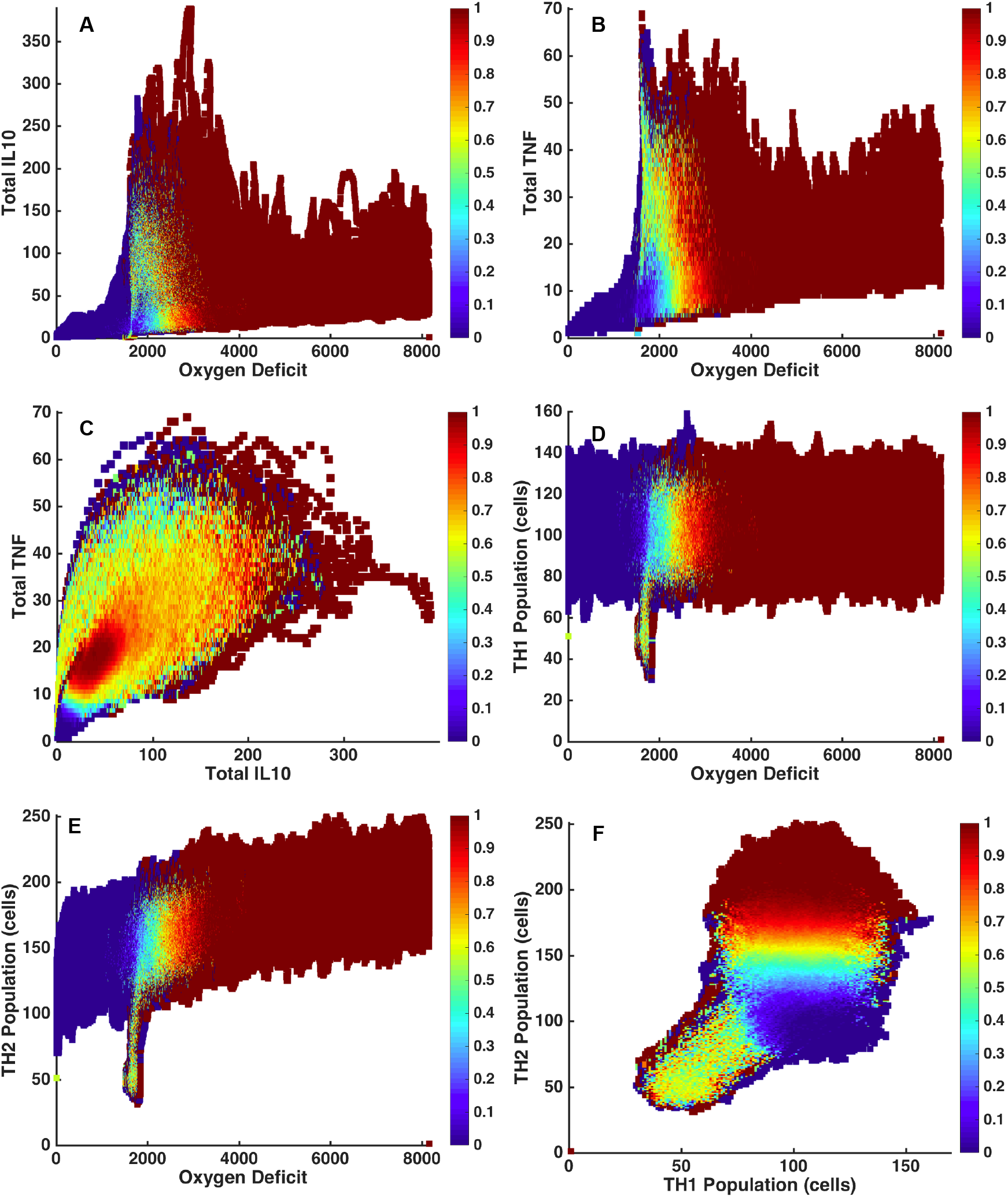
2-dimensional Probabilistic Basins of Attraction (BoA). The PBoA is a heatmap of the death probability for across the system states reachable by the model in parameter space; while this object can be represented as a function of any number of variables measured in the simulation, we present 2-dimensional slices for simplicity and ease of representation/interpretation. The PBoA’s in this figure are generated from the following parameter set: invasiveness=2, toxigenesis=4, environmental toxicity=0, and host resilience=0.1, and have been constructed through the analysis of 1000 trajectories.

### 0.5 Metric #5. Stochastic Trajectory Analysis (STA)

As noted above, biological systems are dynamic systems, in which the behavior of interest is reflected by the trajectory of the system (i.e. disease course). The PBoA provides comprehensive depiction of the predictive capabilities of state space analysis for this model of sepsis; we now turn our attention to providing comprehensive trajectory analysis. As with the other aspects of the investigations presented in this paper, the ability to perform HPC-enabled simulations generates system data at a granularity and scale not feasible in real world systems, with the goal of empirically generating the geometric and topological produced by mathematical functions that can lead to novel, fundamental insights into the system under study. With Stochastic Trajectory Analysis (STA), we examine the specific dynamics associated with the transitions out of the stochastic zone of the PBoA with the goal of more precisely characterizing “tipping points” with respect to system trajectories both at the population and individual level. These simulation experiments are performed with specifically chosen parameter sets and values for initial injury across a set of stochastic replicates (i.e. only differences between runs are the Random Number Seeds). These simulations attempt to characterize the intrinsic stochasticity of the IIRABM and establish boundaries for developing predictive metrics for system outcome. The STA therefore consists of a number of trajectories (N = 1000) plotted with respect to system health, with those trajectories subdivided into 3 zones: 1) inevitable healing, 2) inevitable death and 3) uncertain outcome. Thus the tipping points into inevitable healing and inevitable death are defined for the particular population of stochastic replicates; the robustness of these boundaries are further evaluated by a series of simulations that stop the simulation at or near the boundary, perform re-seeding of the Random Number Generator, and proceed with the simulation run from that point. This accomplishes a means of evaluating the stochastic range around the initially described “tipping boundary,” and serves to provide a representation of trajectory space distribution that corresponds to the stochastic zone of the PBoA. The internal stochasticity of the IIRABM for a particular parameter set is demonstrated Fig. 5 plots 1000 patient trajectories for oxygen deficit (the simulation’s primary metric for patient health) are plotted against time for up to 84 days. All but two of the cases have resolved by this time. The black bars on this plot bound the stochastic region of the simulation – that for which the outcome cannot be definitively predicted. The region of space above the top black bar is under the influence of the Death Attractor and will certainly die. The region of space below the lower black bar is under the influence of the Life Attractor and will certainly heal completely. We should note that these bars are not definitive and their convergence properties are poorly understood at present. This is because the stochastic boundaries in Fig. 5 are derived from 1000 simulations; one can imagine that there would be some shifting and expansion of these boundaries if the number of simulations were increased to 10,000, 100,000, or even 1,000,000, though this would not continue indefinitely – there is 0 probability that a system would be in a position of 99.99% health and spontaneously begin a decay to death. Conversely, there is 0 probability that a system would be in a state of 79.99% damage and spontaneously reverse course and heal completely. The gradient nature of these boundaries are further illustrated in Fig. 6, which displays the health trajectories for a single parameter set with a single initial random number seed (trajectory shown in red). The simulation is then reseeded 100 times (trajectories shown in blue) when the oxygen deficit reaches values of 3000 (Figure 6A), 3500 (Figure 6B), and 4000 (Figure6C). As the simulation moves closer to the Death Attractor, the probability of a positive outcome diminishes. Panel 6A displays a simulation that was reseeded near the center of the stochastic zone; at this point, the outcome far from certain and the primary driver for final outcome is stochastic noise. Panel 6B shows re-seeding when the original simulation was in a more unhealthy state (higher oxygen deficit) and this has significantly skewed the range of outcomes to the point that there is only a 1% chance of survival. At this point, stochastic noise is still an important component of the simulation and has the ability to overpower the influence of the Death Attractor, though it is unlikely. In Panel 6C, the simulation was reseeded at an even unhealthier state; stochastic noise is no longer a relevant simulation for this simulation as death is a certainty. The series of trajectories displayed in Figure 6C show a system that has crossed out of the stochastic zone and into the basin of attraction for the Death Attractor.

**Figure 5.**
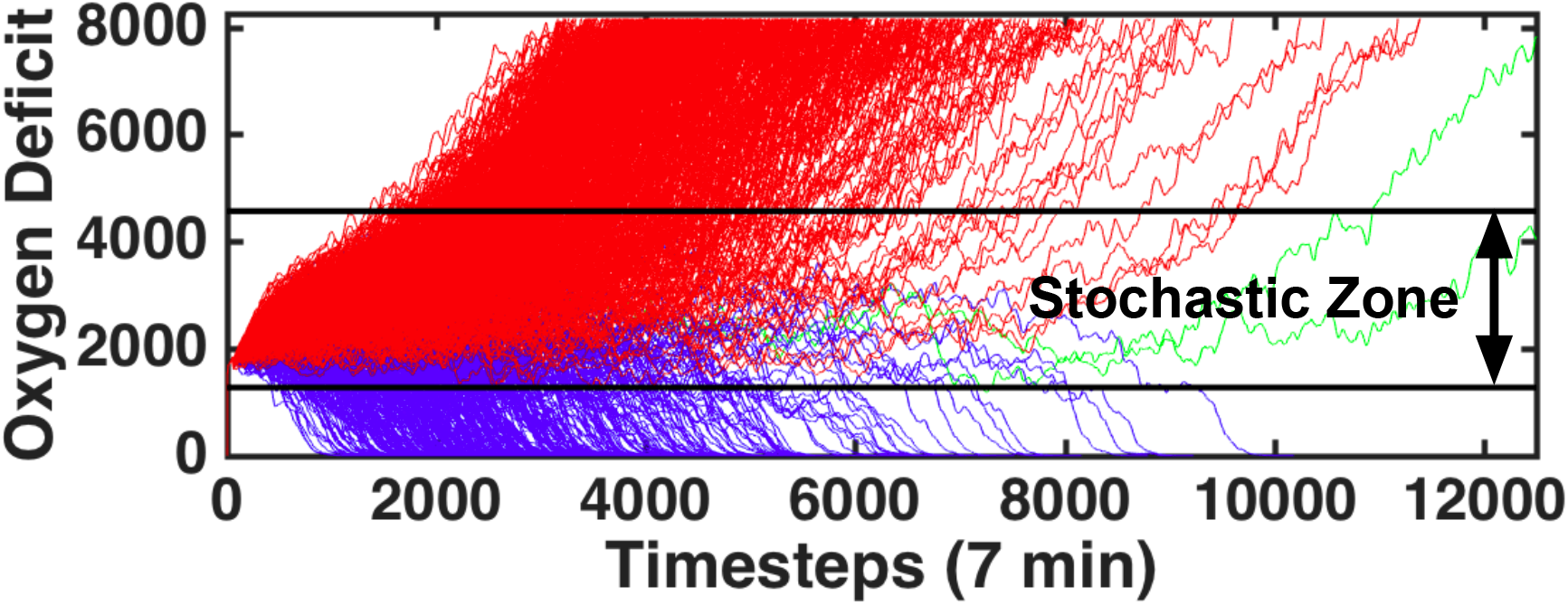
1000 trajectories of the model using the parameter values (invasiveness=2, toxigenesis=4, environmental toxicity=0, and host resilience=0.1) with an injury size of 22. Oxygen Deficit, the simulation’s primary measure of health, is plotted as a function of time. Trajectories colored in red end in death of the patient; trajectories colored in blue end with complete healing of the simulated patient; trajectories in green had not reached a final outcome by 84 simulated days. The black bars represent the boundary of the stochastic zone (the zone in which stochastic noise is (or can be) more powerful than the influence of either attractor). Each bar is a boundary which, once crossed, ultimately determines the simulation’s fate; if an *in silico* patient’s health worsens to the point that it crosses the top black bar, then that patient is sure to die and vice-versa.

**Figure 6.**
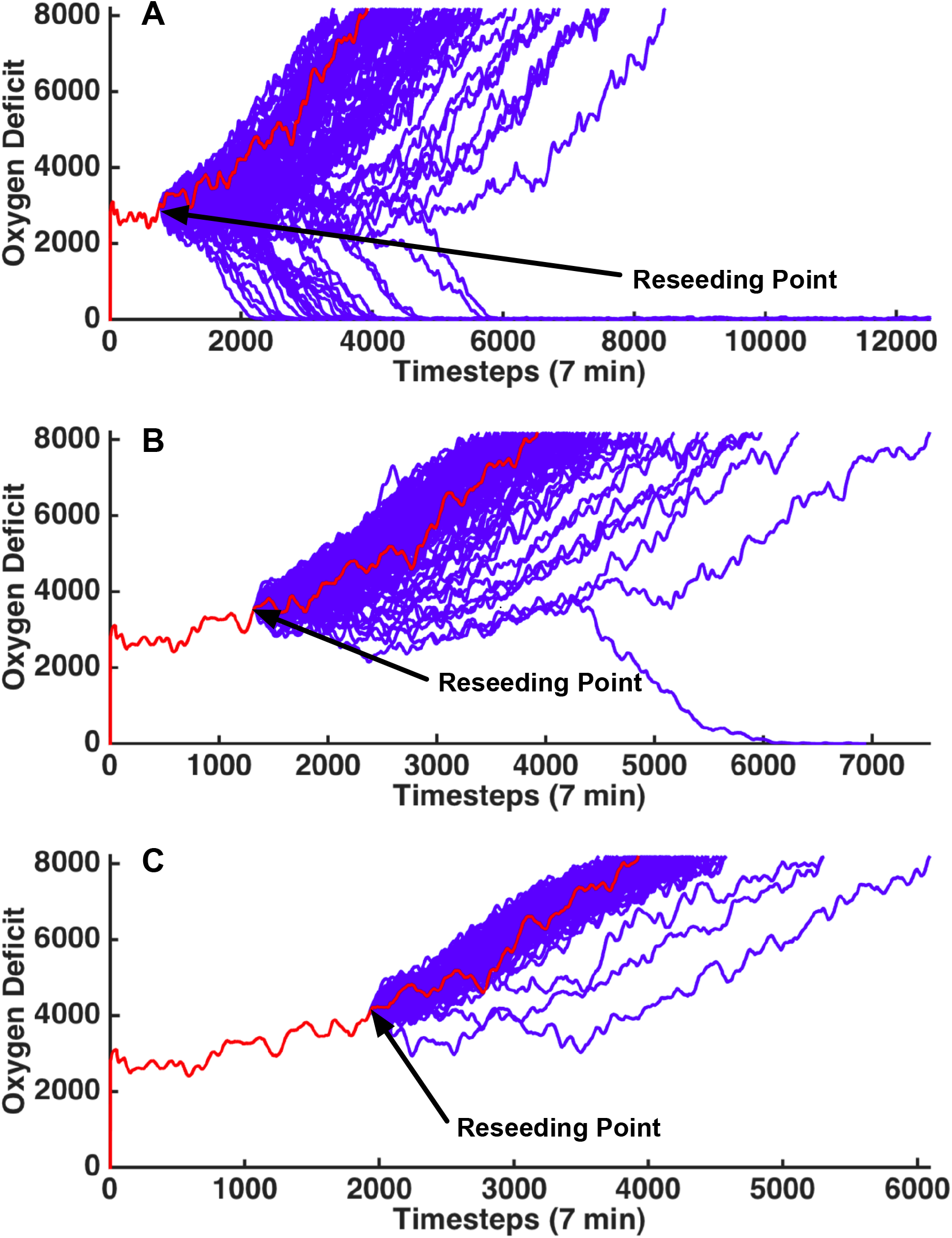
This figure displays patient trajectories for a single parameter set (invasiveness=2, toxigenesis=4, environmental toxicity=0, and host resilience=0.1) and single initial random number seed. The random number generator was re-seeded 100 times at at oxygen deficit=3000 (top), 3500 (middle) and 4000 (bottom). The original trajectory is shown in red and the trajectories generated from reseeding the random number generator are shown in blue. This image further reinforces the identification of a “stochastic zone” (a region of parameter space in which stochastic noise is (or can be) more powerful than the influence of either attractor).

## Discussion

The use of proxy models is ubiquitous and essential to the practice of science. Every laboratory model (in vitro or in vivo) represents a proxy system for the real system being investigated. An essential aspect of all proxy models is that they represent some degree of abstraction or divergence of detail with respect to the real world system; in fact the breadth of their explanatory power is directly related to their generalizability. However, for highly-engineered biological experimental models abstractions/limitations are impossible to characterize formally, with a consequent impact on the ability to extrapolate findings from those models to their reference systems and manifesting in the persistence of the Translational Dilemma [3, 20,21]. To address this dilemma, we propose an alternative approach that uses an abstract, yet sufficiently complex computational proxy model of systemic inflammation/sepsis to establish boundary conditions with respect to the scope of inflammation’s possible behavior. This use of simulation relies on leveraging the benefits of abstraction to increase the generality of conclusions derived through examination of the model, thereby expanding the potential applicability of those results. As opposed to fitting a specific set of experimental/empirical data, we propose utilizing the ability of simulation to vastly increase the “data coverage” of possible states and trajectories of the target system, allowing a more comprehensive representation of the behavior of that system [20,21].

In Ref [22] the authors undertake the near monumental task of compiling the list of randomized clinical trials that form the basis for current critical care practice (see Online Supplement to [22]). However, as daunting (and depressing) as this list is, the sum total of the represented patients, and the data extracted from them, represents only a miniscule fraction and sampling of the possible states and behaviors collected from a model as simplified and abstract as the IIRABM. Visualization of near-comprehensive landscape of the system’s behavior in the current paper illustrates how under-sampling of this landscape can give a skewed view of potential “driving” factors or processes, and lead to overestimation of the general predictive capacity of any algorithm derived from such a sparse sample. The under-sampling problem has been long recognized in the field of signal processing, first described by Nyquist in 1928, and later formalized by Claude Shannon with the Nyquist-Shannon signaling theorem [23, 24], which set lower bounds for a sufficient sampling criterion to discretize a continuous signal. For example, if one were to try to reconstruct the signal *y* = *sin*(*t*) by sampling every *π* seconds starting at *t* = 0, the reconstructed signal would simply be *y* = 0; all of the interesting features of the function would be washed out by under-sampling. While the Nyquist-Shannon theorem deals with 1 dimensional functions we can extrapolate to the current sepsis work, and biomedical research in general. In fact, the “Crisis of Reproducibility” noted in the biomedical arena [25] can be at least partially attributed to the perpetual under-sampling of the total space of possible outcomes, with the consequence that a misleading statistical significance for one particular study will become evident as subsequent studies add to the portions of behavior space sampled [26].

If the sampling is perpetually temporally sparse (as will be the case for the foreseeable future due to the logistical challenges of obtaining good quality clinical data), multiplication of dimensions across which each time point is evaluated only increases the dimensions in which information becomes lost. As we cannot derive a theoretically optimal sampling criterion (the model has no explicit functional form), we instead perform a sufficiently large parameter sweep to bound the region of interest in parameter space – in this specific case that region was the Clinically Relevant region in which any outcome was possible - and provide a reference point by which the degree of under-sampling can be quantified. Without some concept of the overall system behavior space that can approximate the scope of the denominator problem in induction, the danger moving forward is that, given a specific data set, “some” best fit function can be found for that specific data set, but will intrinsically be limited in it’s applicability to the more general condition. This explains why, in terms of clinical decision support as a path towards precision medicine, physiology-based clinical screening tools [27–29] perform essentially as well as multiplexed biomarker/-omics assays [30–33] for their respective predictive targets (onset of sepsis and sepsis outcome). We assert that the perpetual under-sampling of sepsis behavior space places an upper bound on the predictive capacity of any algorithm based on the under-representative data set (which is a pragmatically fixed constraint). The methodology employed for the development of all these scoring systems all assume that the data sets utilized sufficiently represent the overall set of behaviors possible, which the current study demonstrates is not the case. In fact, given the “bowtie structure” of biological signaling and regulatory networks [34], dynamic physiology-based prediction systems [29] [35] [36] likely have a greater chance of success in refining individual patient trajectories, but at the cost of not providing insight into the mechanistic drivers that would be potential targets for therapeutic control. Future work with simulation-based characterization of sepsis will likely involve mapping between computationally-generated behavioral landscapes and finer-grained temporal clinical phenotype characterization using advanced physiological metrics to guide discovery of mechanistic determinants of individual patient trajectories. The importance of identifying general properties of patient trajectories, particularly with respect to the concept of attractors influencing those trajectories, has a long history in the study of sepsis [37,38]. However, moving this recognition beyond the conceptual level has been a challenge over the ensuing decades, due in main part to the fact that the concept of “attractors” derive from dynamical systems theory, which is based on the assumption of deterministic functions (even the study of stochastic dynamical systems primarily involve the addition of a noise term to an existing deterministic function). It is readily clear that biological systems are not deterministic (due to both real and epistemic stochasticity), and are more accurately described as Random Dynamical Systems (RDS) [16]. As noted above, this recognition requires the modification of the classical attractor concept to our Probabilistic Basins of Attraction (PBoA). The concept of the PBoA is predicated on the fact that a “master equation” for the dynamical system is not known; the PBoA can only be derived empirically. And since real-world biological data is too sparse, this requires the use of large-scale simulation experiments to generate the landscapes of the PBoAs, which in turn is only possible with the employment of HPC platforms. In time, for a sufficiently validated model, one can imagine the PBoA as a valuable clinical diagnostic tool. Chemokine levels and their time-derivatives would be used as input coordinates to a high-dimensional matrix containing outcome probabilities. These probabilities would then inform future treatment paths. Since they are derived from mechanistic simulations, PBoAs can also be used to generate targets and hypotheses for therapy discovery, e.g., if it were shown that the death outcome is strongly associated with a specific signal configuration, this would suggest control strategies aimed at that particular set of signals, subsequently evaluated by examining the change to the distribution of the PBoA. Additionally, Stochastic Trajectory Analysis (STA) can further parse the dynamics and distribution of the trajectory space in the transition zones between stochastic indeterminant behavior and movement towards the probabilistic attractors. This will help guide the “right time” component of Precision Medicine that is all too often overlooked, as well help define the generalizability of putative control strategies by evaluation across STAs for individual parameter conditions (i.e. patient types). We have already begun investigation of the use of deep reinforcement learning and evolutionary computing/genetic programming to search across the space of possible interventions/control strategies guided by PBoAs and STAs to first determine controllability of the overall system, and subsequently attempt to determine the robustness/scalability of control strategies/policies identified. We propose that proxy simulation models, such as the IIRABM, can be refined with an iterative loop between simulation and empiric plausibility evaluation (just as meteorological simulations are constantly being refined), and during this process can be used to help establish boundaries for plausible and implausible investigatory strategies, define expectations for possible success and potentially eliminate futile approaches [41]. Absent this approach, currently, prospective analysis of putative stratification systems (to say nothing of potential therapeutic interventions) cannot be done from a pure real-world empirical, data-centric fashion: such failures will only become evident ex post facto, with the consequent loss of time, money, resources and lives [42–50]. Akin to the current use of HPC resources in physics and meteorology, the approach demonstrated in this paper, the use of extremely large scale simulation and simulated data, provides the only demonstrated effective scientific strategy to prospectively identify the boundaries of fruitful investigation. The use of large-scale simulation based science for sepsis specifically, and biomedicine in general, represents a potential path towards the full-scale application of engineering and control principles to the care of individuals in terms of personalized/precision medicine and truly rational design of effective therapeutics.

## Acknowledgments and Conflicts of Interest

Drs. An and Cockrell are supported by high performance computing resources of the National Energy Research Scientific Computing Center, a DOE Office of Science User Facility supported by the Office of Science of the U.S. Department of Energy under Contract No. DE-AC02-05CH1123, and by computing resources provided by the University of Chicago Computation Institute (Beagle2). Both Drs. An and Cockrell are supported under grant 1S10OD018495-01 from the National Institutes of Health, as well as by funds from Lawrence Livermore National Laboratory under Award #B616283. Dr. An is also a paid consultant for Immunetrics, Inc., but that role is unrelated to the content of this manuscript.

## References

1. Wood KA, Angus DC: Pharmacoeconomic implications of new therapies in sepsis. Pharmacoeconomics 2004, 22(14):895–906.

2. Angus DC, Barnato AE, Bell D, Bellomo R, Chong CR, Coats TJ, Davies A, Delaney A, Harrison DA, Holdgate A et al: A systematic review and meta-analysis of early goal-directed therapy for septic shock: the ARISE, ProCESS and ProMISe Investigators. Intensive Care Med 2015, 41(9):1549–1560.

3. An G: Closing the scientific loop: bridging correlation and causality in the petaflop age. Science translational medicine 2010, 2(41):41ps34.

4. Jonas E, Kording KP: Could a Neuroscientist Understand a Microprocessor? PLoS Comput Biol 2017, 13(1):e1005268.

5. An G, Fitzpatrick BG, Christley S, Federico P, Kanarek A, Neilan RM, Oremland M, Salinas R, Laubenbacher R, Lenhart S: Optimization and Control of Agent-Based Models in Biology: A Perspective. Bull Math Biol 2016.

6. An G: In silico experiments of existing and hypothetical cytokine-directed clinical trials using agent-based modeling. Critical care medicine 2004, 32(10):2050–2060.

7. Tamayo E, Fernandez A, Almansa R, Carrasco E, Heredia M, Lajo C, Goncalves L, Gomez-Herreras JI, de Lejarazu RO, Bermejo-Martin JF: Pro- and anti-inflammatory responses are regulated simultaneously from the first moments of septic shock. Eur Cytokine Netw 2011, 22(2):82–87.

8. Osuchowski MF, Welch K, Siddiqui J, Remick DG: Circulating cytokine/inhibitor profiles reshape the understanding of the SIRS/CARS continuum in sepsis and predict mortality. J Immunol 2006, 177(3):1967–1974.

9. Hotchkiss RS, Monneret G, Payen D: Immunosuppression in sepsis: a novel understanding of the disorder and a new therapeutic approach. Lancet Infect Dis 2013, 13(3):260–268.

10. Hotchkiss RS, Monneret G, Payen D: Sepsis-induced immunosuppression: from cellular dysfunctions to immunotherapy. Nat Rev Immunol 2013, 13(12):862–874.

11. Boomer JS, To K, Chang KC, Takasu O, Osborne DF, Walton AH, Bricker TL, Jarman SD, 2nd, Kreisel D, Krupnick AS et al: Immunosuppression in patients who die of sepsis and multiple organ failure. JAMA 2011, 306(23):2594–2605.

12. Wilensky U, Evanston I: NetLogo: Center for connected learning and computer-based modeling. Northwestern University, Evanston, IL 1999:49–52.

13. An G: Agent-based computer simulation and sirs: building a bridge between basic science and clinical trials. Shock 2001, 16(4):266–273.

14. Strogatz SH: Nonlinear dynamics and chaos: with applications to physics, biology, chemistry, and engineering: Westview press; 2014.

15. Attractor [http://mathworld.wolfram.com/Attractor.html]

16. Arnold L: Random dynamical systems: Springer Science & Business Media; 2013.

17. Ferguson N, Galley H, Webster N: T helper cell subset ratios in patients with severe sepsis. Intensive care medicine 1999, 25(1):106–109.

18. Biron BM, Ayala A, Lomas-Neira JL: Biomarkers for Sepsis: What Is and What Might Be? Biomark Insights 2015, 10(Suppl 4):7–17.

19. Faix JD: Biomarkers of sepsis. Crit Rev Clin Lab Sci 2013, 50(1):23–36.

20. An G: Small to large, lots to some, many to few: in silico navigation of the Translational Dilemma. Crit Care Med 2012, 40(4):1334–1335.

21. An G, Christley S: Addressing the translational dilemma: dynamic knowledge representation of inflammation using agent-based modeling. Crit Rev Biomed Eng 2012, 40(4):323–340.

22. Buchman TG, Billiar TR, Elster E, Kirk AD, Rimawi RH, Vodovotz Y, Zehnbauer BA: Precision Medicine for Critical Illness and Injury. Crit Care Med 2016, 44(9):1635–1638.

23. Shannon C: A mathematical theory of communication, Bell System Technical Journal. In: Mathematical Reviews. vol. MR10. MathSciNet; 1948:133e.

24. Black H: Modulation Theory. NY and Toronto: D van Nostrand Co; 1953.

25. Baker M: 1,500 scientists lift the lid on reproducibility. Nature 2016, 533(7604):452–454.

26. Szucs D, Ioannidis JP: Empirical assessment of published effect sizes and power in the recent cognitive neuroscience and psychology literature. PLoS Biol 2017, 15(3):e2000797.

27. Moore LJ, Jones SL, Kreiner LA, McKinley B, Sucher JF, Todd SR, Turner KL, Valdivia A, Moore FA: Validation of a screening tool for the early identification of sepsis. J Trauma 2009, 66(6):1539–1546; discussion 1546-1537.

28. Wawrose R, Baraniuk M, Standiford L, Wade C, Holcomb J, Moore L: Comparison of Sepsis Screening Tools’ Ability to Detect Sepsis Accurately. Surg Infect (Larchmt) 2016, 17(5):525–529.

29. Lindner HA, Balaban U, Sturm T, Wei C, Thiel M, Schneider-Lindner V: An Algorithm for Systemic Inflammatory Response Syndrome Criteria-Based Prediction of Sepsis in a Polytrauma Cohort. Crit Care Med 2016, 44(12):2199–2207.

30. Bauer M, Giamarellos-Bourboulis EJ, Kortgen A, Moller E, Felsmann K, Cavaillon JM, Guntinas-Lichius O, Rutschmann O, Ruryk A, Kohl M et al: A Transcriptomic Biomarker to Quantify Systemic Inflammation in Sepsis - A Prospective Multicenter Phase II Diagnostic Study. EBioMedicine 2016, 6:114–125.

31. Mickiewicz B, Vogel HJ, Wong HR, Winston BW: Metabolomics as a novel approach for early diagnosis of pediatric septic shock and its mortality. Am J Respir Crit Care Med 2013, 187(9):967–976.

32. Langley RJ, Tsalik EL, van Velkinburgh JC, Glickman SW, Rice BJ, Wang C, Chen B, Carin L, Suarez A, Mohney RP et al: An integrated clinico-metabolomic model improves prediction of death in sepsis. Science translational medicine 2013, 5(195):195ra195.

33. Neugebauer S, Giamarellos-Bourboulis EJ, Pelekanou A, Marioli A, Baziaka F, Tsangaris I, Bauer M, Kiehntopf M: Metabolite Profiles in Sepsis: Developing Prognostic Tools Based on the Type of Infection. Crit Care Med 2016, 44(9):1649–1662.

34. Doyle J, Csete M: Motifs, control, and stability. PLoS Biol 2005, 3(11):e392.

35. Moorman JR, Lake DE, Ivanov P: Early Detection of Sepsis-A Role for Network Physiology? Crit Care Med 2016, 44(5):e312–313.

36. Moss TJ, Lake DE, Calland JF, Enfield KB, Delos JB, Fairchild KD, Moorman JR: Signatures of Subacute Potentially Catastrophic Illness in the ICU: Model Development and Validation. Crit Care Med 2016, 44(9):1639–1648.

37. Buchman TG: Physiologic stability and physiologic state. The Journal of trauma 1996, 41(4):599–605.

38. Godin PJ, Buchman TG: Uncoupling of biological oscillators: a complementary hypothesis concerning the pathogenesis of multiple organ dysfunction syndrome. Critical care medicine 1996, 24(7):1107–1116.

39. Cohen M, Falcone P, Kusnezov D, Paragas J: Sepsis and Presidential Initiatives. Crit Care Med 2016, 44(11):1963–1965.

40. Brown D, Namas RA, Almahmoud K, Zaaqoq A, Sarkar J, Barclay DA, Yin J, Ghuma A, Abboud A, Constantine G et al: Trauma in silico: Individual-specific mathematical models and virtual clinical populations. Science translational medicine 2015, 7(285):285ra261.

41. Vodovotz Y, An G: A Roadmap for a Rational Future: A Systematic Path for the Design and Implementation of New Therapeutics. In: Translational Systems Biology: Concepts and Practice for the Future of Biomedical Research. edn. Waltham, MA: Elsevier; 2014: 69–78.

42. Reinhart K, Menges T, Gardlund B, Zwaveling JH, Smithes M, Vincent J-L, Tellado JM, Salgado-Remigio A, Zimlichman R, Withington S: Randomized, placebo-controlled trial of the anti-tumor necrosis factor antibody fragment afelimomab in hyperinflammatory response during severe sepsis: The RAMSES Study. Critical care medicine 2001, 29(4):765–769.

43. Opal SM, Fisher CJ, Dhainaut J-FA, Vincent J-L, Brase R, Lowry SF, Sadoff JC, Slotman GJ, Levy H, Balk RA: Confirmatory interleukin-1 receptor antagonist trial in severe sepsis: a phase III, randomized, doubleblind, placebo-controlled, multicenter trial. Critical care medicine 1997, 25(7):1115–1124.

44. Rhee P, Morris J, Durham R, Hauser C, Cipolle M, Wilson R, Luchette F, McSwain N, Miller R: Recombinant humanized monoclonal antibody against CD18 (rhu MAb CD18) in traumatic hemorrhagic shock: Results of a phase II clinical trial. Journal of Trauma and Acute Care Surgery 2000, 49(4):611–620.

45. Fisher CJ, Dhainaut J-FA, Opal SM, Pribble JP, Balk RA, Slotman GJ, Iberti TJ, Rackow EC, Shapiro MJ, Greenman RL: Recombinant human interleukin 1 receptor antagonist in the treatment of patients with sepsis syndrome: results from a randomized, double-blind, placebo-controlled trial. Jama 1994, 271(23):1836–1843.

46. Fisher Jr CJ, Agosti JM, Opal SM, Lowry SF, Balk RA, Sadoff JC, Abraham E, Schein RM, Benjamin E: Treatment of septic shock with the tumor necrosis factor receptor: Fc fusion protein. New England Journal of Medicine 1996, 334(26):1697–1702.

47. Abraham E, Glauser MP, Butler T, Garbino J, Gelmont D, Laterre PF, Kudsk K, Bruining HA, Otto C, Tobin E: p55 tumor necrosis factor receptor fusion protein in the treatment of patients with severe sepsis and septic shock: A randomized controlled multicenter trial. Jama 1997, 277(19):1531–1538.

48. Abraham E, Anzueto A, Gutierrez G, Tessler S, San Pedro G, Wunderink R, Dal Nogare A, Nasraway S, Berman S, Cooney R: Double-blind randomised controlled trial of monoclonal antibody to human tumour necrosis factor in treatment of septic shock. The Lancet 1998, 351(9107):929–933.

49. Abraham E, Laterre P-F, Garbino J, Pingleton S, Butler T, Dugernier T, Margolis B, Kudsk K, Zimmerli W, Anderson P: Lenercept (p55 tumor necrosis factor receptor fusion protein) in severe sepsis and early septic shock: a randomized, double-blind, placebo-controlled, multicenter phase III trial with 1,342 patients. CRITICAL CARE MEDICINE-BALTIMORE-2001, 29(3):503–510.

50. Root RK, Lodato RF, Patrick W, Cade JF, Fotheringham N, Milwee S, Vincent J-L, Torres A, Rello J, Nelson S: Multicenter, double-blind, placebo-controlled study of the use of filgrastim in patients hospitalized with pneumonia and severe sepsis. Critical care medicine 2003, 31(2):367–373.

